# Structural Pathway for Allosteric Activation of the Autophagic PI 3-Kinase Complex I

**DOI:** 10.1101/758797

**Authors:** Lindsey N. Young, Felix Goerdeler, James H. Hurley

## Abstract

Autophagy induction by starvation and stress involves the enzymatic activation of the class III phosphatidylinositol 3-kinase complex I (PI3KC3-C1). The inactive basal state of PI3KC3-C1 is maintained by inhibitory contacts between the VPS15 protein kinase and VPS34 lipid kinase domains that restrict the conformation of the VPS34 activation loop. Here, the pro-autophagic MIT domain-containing protein NRBF2 was used to map the structural changes leading to activation. Cryo-EM was used to visualize stepwise PI3KC3-C1 activating effects of binding the NRFB2 MIT domains. Binding of a single NRBF2 MIT domain to bends the helical solenoid of the VPS15 scaffold, displaces the protein kinase domain of VPS15, and releases the VPS34 kinase domain from the inhibited conformation. Binding of a second MIT stabilizes the VPS34 lipid kinase domain in an active conformation that has an unrestricted activation loop and is poised for access to membranes.

---

Autophagy is a core cellular process, conserved throughout eukaryotes, which is the central recycling system for the removal of misfolded proteins, damaged organelles, and the recycling of nutrients in starvation. Autophagic dysfunction is implicated in many disease states, including neurodegeneration, immune disorders, cancer, and aging, among others (1). The class III phosphatidylinositol-3 kinase complexes (PI3KC3) I and II (PI3KC3-C1 and -C2), respectively, are essential for the initiation and expansion of autophagosomes (2–5). PI3KC3 generates the lipid phosphatidylinositol-3-phosphate, PI(3)P, which is recognized by the WIPI proteins. WIPIs in turn recruit the machinery that conjugates the autophagosomal marker LC3 to the expanding autophagosomal membrane (6). PI3KC3-C1 has been proposed to be a promising therapeutic target for autophagy activators (7) because the generation of PI(3)P is absolutely required for the recruitment of downstream autophagy proteins. There is considerable medical interest in selectively activating this pathway to promote human health and treat disease, yet there are no FDA approved pharmaceuticals that uniquely activate autophagy.

PI3KC3-C1 consists of the lipid kinase VPS34, the putative serine/threonine protein kinase VPS15, the regulatory subunit BECN1, and the early autophagy-specific targeting subunit ATG14 (8, 9). In PI3KC3-C2, ATG14 is replaced with UVRAG (10), while the other three subunits are preserved. The overall architecture of both PI3KC3-C1 and -C2 has the shape of the letter V (11, 12) (Fig. 1A). The long coiled-coils of BECN1 and ATG14 scaffold the left arm in the standard view, with the membrane binding BARA domain of BECN1 located at the outermost tip of the arm. PI3KC3-C2 is inhibited by Rubicon and the HIV-1 protein Nef (13–15), which regulate membrane docking by the tip of the left arm. The catalytic domains of the kinases VPS34 and VPS15 are at the tip of the right arm (11, 12). PI3KC3 complexes are phosphoregulated by the Unc-51 like autophagy-activating kinase 1 (ULK1) complex (16), among other kinases (3). PI3KC3-C1 is inhibited by anti-apoptotic proteins Bcl-2/Bcl-XL (10). The binding of Bcl-2/Bcl-XL and the best-characterized phosphoregulatory modulations of PI3KC3-C1 map to the base of the V. Thus far, the structural mechanisms of the many PI3KC3 regulators acting at the base of the V have been undefined.

**Fig. 1.**
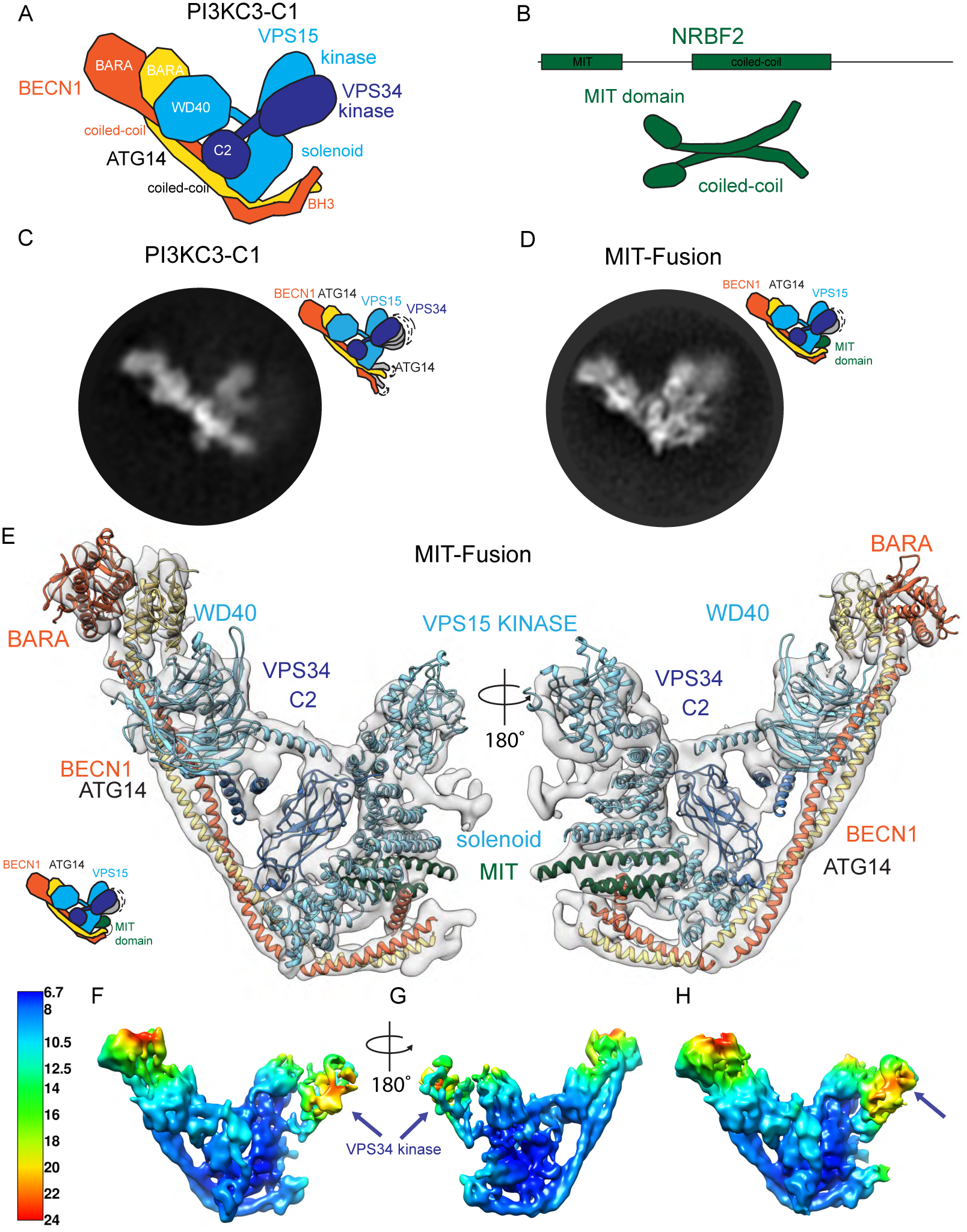
CryoEM structure of “MIT-Fusion” (BECN1-NRBF2^MIT^ PI3KC3-C1). (A) Schematic of PI3KC3-C1 containing VPS34, VPS15, BECN1, and ATG14. (B) Schematic of NRBF2 (C) 2D cryoEM class average of PI3KC3-C1 for comparison, illustrating the mobility of the VPS34 kinase domain. (D) 2D cryoEM class average of MIT-Fusion. (E) CryoEM reconstruction of MIT-Fusion with VPS34 helical and catalytic domains masked out. Models originate from PDB depositions for yeast PI3KC3-C2 (5DFZ), NRBF2^MIT^ (4ZEY), and BECN1^BARA^ (4DDP). The cryoEM map has been deposited in the EMDB under 20387. (F) Local resolution of MIT-Fusion structure with VPS34 helical and kinase domains included in masking, shown at a contour level of 10 σ or 0.0097 in Chimera and rotated 180° in (G). (H) The VPS34 catalytic domain is visible (arrow), albeit at a low resolution of 18-24 Å, when contoured at a threshold of 6 σ or 0.0058 Chimera.

Nuclear Receptor Binding Factor 2 (NRBF2) is a positive regulator of PI3KC3-C1 that also acts at the base of the V shape (17). NRBF2 was identified as a PI3KC3-C1 associated factor through proteomics of the mammalian autophagy network (18). Depletion of NRBF2 reduced autophagosome formation, implicating NRBF2 as pro-autophagic (18). Work in *S. cerevisiae* identified Atg38 as the homolog of NRBF2 and confirmed Atg38 to be a pro-autophagic component of PI3KC3-C1 (19). The pro-autophagic function of NRBF2 has been confirmed by most reports in mammalian cells (20–24), although contrary findings were reported by one group (24). NRBF2 and its ortholog Atg38 contain an N-terminal Microtubule Interacting and Trafficking (MIT) domain that binds to PI3KC3-C1, followed by a central dimeric coiled-coil domain (17, 19, 25) (Fig. 1B). Dimerization can be decoupled from activation, as the N-terminal MIT domain (NRBF2^MIT^) alone is sufficient to enhance the kinase activity of the autophagy specific PI3-Kinase *in vitro* (17) and to rescue autophagy induction in MEFs (21). Here, we used cryo-EM and allied methods to show how NRBF2 allosterically activates PI3KC3-C1 at a structural and mechanistic level.

## Results

### Characterization of PI3KC3-C1 containing a NRBF2-MIT-BECN1 fusion construct

We had previously found that (1) the isolated NRBF2^MIT^ was sufficient to activate PI3KC3-C1, and (2) that the principal binding site of NRBF2^MIT^ included the N-terminal domain of BECN1 (17). In order to generate a non-dissociable NRBF2^MIT^ complex with PI3KC3-C1 for cryo-EM studies, we built on these observations by fusing NRBF2^MIT^ to the N-terminus of BECN1 using a flexible (Gly-Gly-Ser)_4_ linker (Fig. S1A, B). To confirm that the MIT-linker construct occupies the same binding site within PI3KC3-C1 as unfused NRBF2, we purified a variant PI3KC3-C1 in which wild-type BECN1 was replaced by NRBF2^MIT^-BECN1 (“MIT-Fusion”), performed HDX-MS, and compared the HDX protection of the NRBF2 complex to the fusion. In both cases, the reference was taken as wild-type PI3KC3-C1 in the absence of NRBF2 (Fig. S1C).

We observed protection patterns in VPS34 (Fig. S1D), BECN1 (Fig. S1E), ATG14 (Fig. S1F), and VPS15 (Fig. S1G) consistent with NRBF2 binding near the base of the complex. Peptides within the N-terminal domains of BECN1 and ATG14, the VPS15 helical solenoid, and the VPS34 C2 domain showed HDX decreases of up to 50%. However, most HDX decreases throughout the remainder of the four subunits did not exceed ∼10 %. In the NRBF2-BECN1 fusion complex, the regions that showed the highest protection were typically even more protected (Fig S1D-G). Two regions of ATG14 that were ∼15% protected in the NRBF2 complex with ∼30% protected in the fusion, for example (Fig. S1F). Therefore, the presence of the fused NRBF2^MIT^ generally recapitulates the qualitative protection pattern seen in the non-covalent complex with intact NRBF2, but the degree of protection is greater. This is presumably because the fused complex cannot dissociate when it is diluted into D_2_O for exchange experiments. Outside of the regions expected to interact with NRBF2, increased protection was also noted. For example, most of the catalytic domains of VPS34 and VPS15 manifested a ∼20% increase in protection in the fusion (Fig. S1D, G). These increases are suggestive of a more global decrease in protein dynamics in the fused construct.

### Cryo-EM structure of the BECN1-MIT-Fusion Complex

Previous EM analyses of PI3KC3s have been limited by the dynamic character of these complexes and by their tendency to be in preferred orientations on EM grids (11, 17, 25, 26). Even when preferred orientations can be reduced or eliminated, the dynamics of the catalytic right arm of the complex (27) has still limited analysis (14). The apparent reduction in dynamics in the BECN1-NRBF2^MIT^ form of PI3KC3-C1 suggested that this sample might be more tractable to cryo-EM than the wild-type PI3KC3-C1 and -C2 complexes. Two-dimensional class averages of BECN1-NRBF2^MIT^ PI3KC3-C1 showed additional features compared to wild-type PI3KC3-C1 (Fig. 1C, D). A reconstruction was obtained with an overall resolution of 7.7 Å and local resolution ranging from 6.7-24 Å (Fig. 1E, F, Fig. S2, Table S1), showing the expected V-shaped architecture. Atomic coordinates were modeled taking the crystal structures of yeast PI3KC3-C2 (12) and the NRBF2^MIT^ (RCSB entry 4ZEY) as the starting point (Fig. 1E). Density for the catalytic domains at the tip of the right arm is less defined than for the rest of the structure, however the density was clear enough to show that the VPS15 kinase is displaced relative to its position in the yeast PI3KC3-C2 structure. Despite the lack of side-chain definition at the attained resolution, it was possible to dock the NRBF2^MIT^ into density unambiguously on the basis of the unequal lengths of the three MIT helices.

The NRBF2 MIT is localized near the base of the right arm, on the back side of the complex in the standard view (Fig. 1E). The extended contacts seen with multiple subunits are consistent with the high affinity (40 nM) of NRBF2 for PI3KC3-C1 (17). The MIT interacts extensively with the central part of the VPS15 helical solenoid. The MIT also interacts with a helical density associated with the N-terminal portions of BECN1 and ATG14 (Fig. 1E). On the basis of its high HDX protection (Fig. S1E), this helix was provisionally assigned as the BH3 domain of BECN1. This region has not previously been visualized in any of the PI3KC3 structures, and it therefore appears to become ordered only in the presence of NRBF2^MIT^. This helix in turn interacts with the most N-terminal portions of the parallel BECN1 and ATG14 coiled coils (Fig. 1E).

### Activation requires NRBF2^MIT^ binding at two sites

The isolated NRBF2^MIT^ domain, added at a 60-fold molar excess to PI3KC3-C1, enhanced lipid kinase activity to the same degree as the full-length dimeric NRBF2 (Fig. 2A) (17). Given that observation and the apparent full occupancy of the MIT binding site in the cryo-EM structure, we had expected that the BECN1-NRBF2^MIT^ fusion C1 complex would also have enhanced activity. Contrary to expectations, the BECN1-NRBF2^MIT^ fusion C1 complex had essentially the same activity as the C1 complex in the absence of NRBF2 (Fig. 2A). Addition of NRBF2^MIT^ to the BECN1-NRBF2^MIT^ fusion C1 complex enhanced activity to a similar extent as for the wild-type complex (Fig. 2A). We interpret this to mean that activation of PI3KC3-C1 by NRBF2 requires NRBF2^MIT^ to engage with at least two distinct binding sites on PI3KC3-C1.

**Fig. 2.**
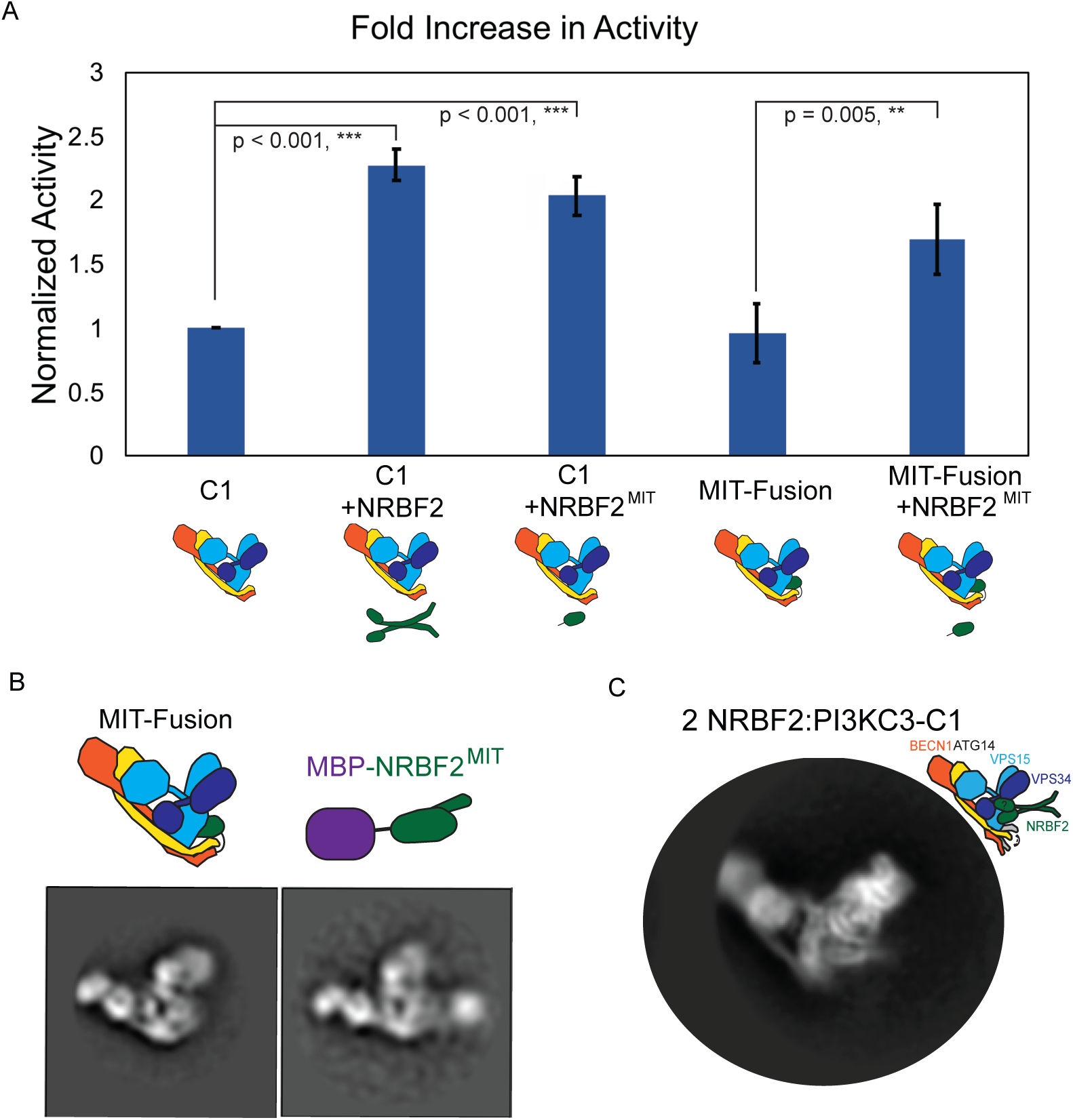
Two copies NRBF2^MIT^ are required for activation. (A) Activity assay of PI3KC3-C1 and MIT-Fusion, with or without NRBF2^MIT^ or NRBF2. The average of three experiments performed in duplicate and normalized to PI3KC3-C1 activity are shown. The error bars represent the standard error of the mean and p-values were determined between PI3KC3-C1 and NRBF2^MIT^ or NRBF2, and MIT-Fusion and NRBF2. (B) Negative stain 2D class average of MIT-Fusion and MIT-Fusion incubated with MBP-NRBF2^MIT^. (C) 2D cryoEM class average of full length NRBF2 bound to PI3KC3-C1.

In order to directly test the existence of a second MIT binding site, negative stain (NS) EM was carried out for the BECN1-NRBF2^MIT^ fusion C1 in the presence of a maltose binding protein (MBP) -tagged-NRBF2^MIT^ construct. Two-dimensional class averages of NS-EM images showed that a second copy of NRBF2^MIT^ binds to the MIT-Fusion complex as shown by the extra density for the MBP-tagged-NRBF2^MIT^ (Fig. 2B). This extra density is located at the base of the complex near the first binding site, suggesting that both N-terminal MIT domains of a single full-length dimeric NRBF2 can occupy both sites simultaneously. These data suggest that the second binding site is lower affinity than the first, and that both binding sites are required for enzymatic activation.

### Cryo-EM structure of full-length NRBF2 bound to PI3KC3-C1

A cryo-EM dataset was obtained for PI3KC3-C1 bound to full-length NRBF2. Two-dimensional class averages showed additional ordered features at the base of the complex and on the right arm as compared to the MIT fusion (Fig. 2C). Single particle reconstruction was carried out to an overall resolution of 6.6 Å (Fig. 3A-E, S3, Table S1), with local resolution ranging from 5.4 to 18 Å (Fig. 3F). The position of the first NRBF2^MIT^ is essentially identical to that seen in the MIT fusion. The local resolution of the NRBF2^MIT^ is improved, enabling the MIT three-helix bundle to be placed with greater precision (Fig. 3C-E). The structure shows the N-terminus of MIT α1, the α2-α3 connector, and the length of α3 of NRBF2^MIT^ with helices α12, α14, and α16 of the VPS15 solenoid (Fig. 3E).

**Fig. 3.**
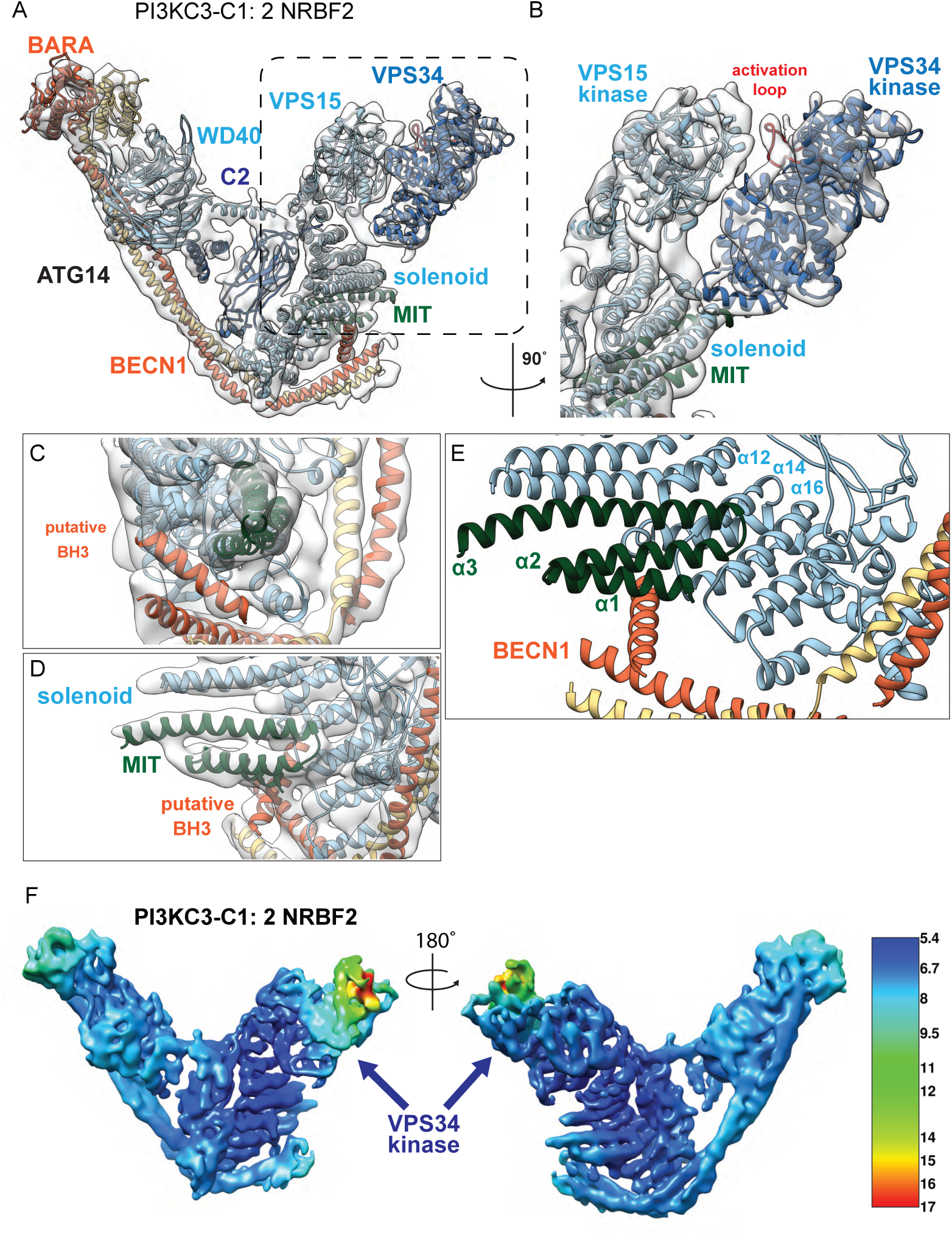
CryoEM structure containing PI3KC3-C1 and full length, dimeric NRBF2. (A) CryoEM reconstruction of a sample containing PI3KC3-C1 and full length NRBF2 (PI3KC3-C1:2NRBF2) contoured at 10 σ or 0.0144 in Chimera. The cryoEM map has been deposited in the EMDB under 20390. Models docked into cryoEM reconstruction include yeast PI3KC3-C2 (5DFZ), VPS34 kinase domain (4PH4), NRBF2^MIT^ (4ZEY), and BECN1^BARA^ (4DDP). (B) Close-up of the cryoEM reconstruction of the VPS15 kinase (cyan, PDB: 5DFZ) and VPS34 kinase domains (blue, PDB: 4PH4), activation loop of VPS34 in red. (C) Close-up of the cryoEM reconstruction for the regions contacting NRBF2^MIT^ (red) include density for a helix, putatively, the BH3 helix of BECN1 (orange). (D) Close-up of solenoid of VPS15 (cyan) which provides an extensive face for MIT binding. (E) Cartoon representation of α-helices-1, 2, 3 of NRBF2^MIT^ contacting α-helices 12, 14, 16 of the VPS15 solenoid as well as the putative BH3 helix. (F) Local resolution estimates for PI3KC3:2NRBF2, with the VPS34 catalytic domain defined at 8-18 Å resolution (arrow).

The VPS34 kinase domain in the NRBF2 complex is better ordered than in the MIT fusion, the main point of difference between the two. The VPS34 kinase is still the least well-defined part of the structure, however, as measured by local resolution (Fig. 3F). Although most of the kinase domain is at ∼8 Å resolution (Fig. 3F), the most distal portions are as low as 18 Å resolution. Despite indications of additional features and what appears to be the coiled-coil stalk of dimeric NRBF2 projecting away from the right arm in the 2D class averages (Fig. 2C), it was not possible to assign the locations of these moieties definitively to the density features at the periphery of the VPS34 kinase domain. We infer that the second NRBF2^MIT^ is bound to this relatively flexible region, and it is not surprising that small domains associated with mobile portions of a complex would not be evident.

### Comparison to inactive conformation of PI3KC3-C2

In the yeast crystal structure, the VPS15 kinase domain contacts the activation loop of VPS34 such that VPS15 inhibits basal VPS34 activity (12) (Fig. 4A-C). Binding of the first NRBF2^MIT^ to the VPS15 solenoid alters its bend such that changes are propagated to the N-terminus of the solenoid where it meets the VPS15 kinase domain (Fig. 4A, B). This change pushes the N-terminal part of the solenoid in the outward direction relative to the base of the V. This in turn pushes the VPS15 kinase away from the VPS34 kinase by ∼10 Å near to solenoid and ∼20 Å at the tip (Fig. 4B). This movement of VPS15 breaks the inhibitory contacts with the VPS34 activation loop (Fig. 4D). This triggers the final change in the series, a 25° pivot of VPS34 about the base of the kinase domain such that the distal tip of the kinase domain moves 45 Å (Fig. 4D). These changes separate the two domains so that the gap between the most C-terminal ordered residue of VPS34 and the N-terminal residue of VPS15 increases from 28 to 51 Å (Fig. 4C, D). This change completely liberates the VPS34 kinase activation loop seen in the inactive structure.

**Fig. 4.**
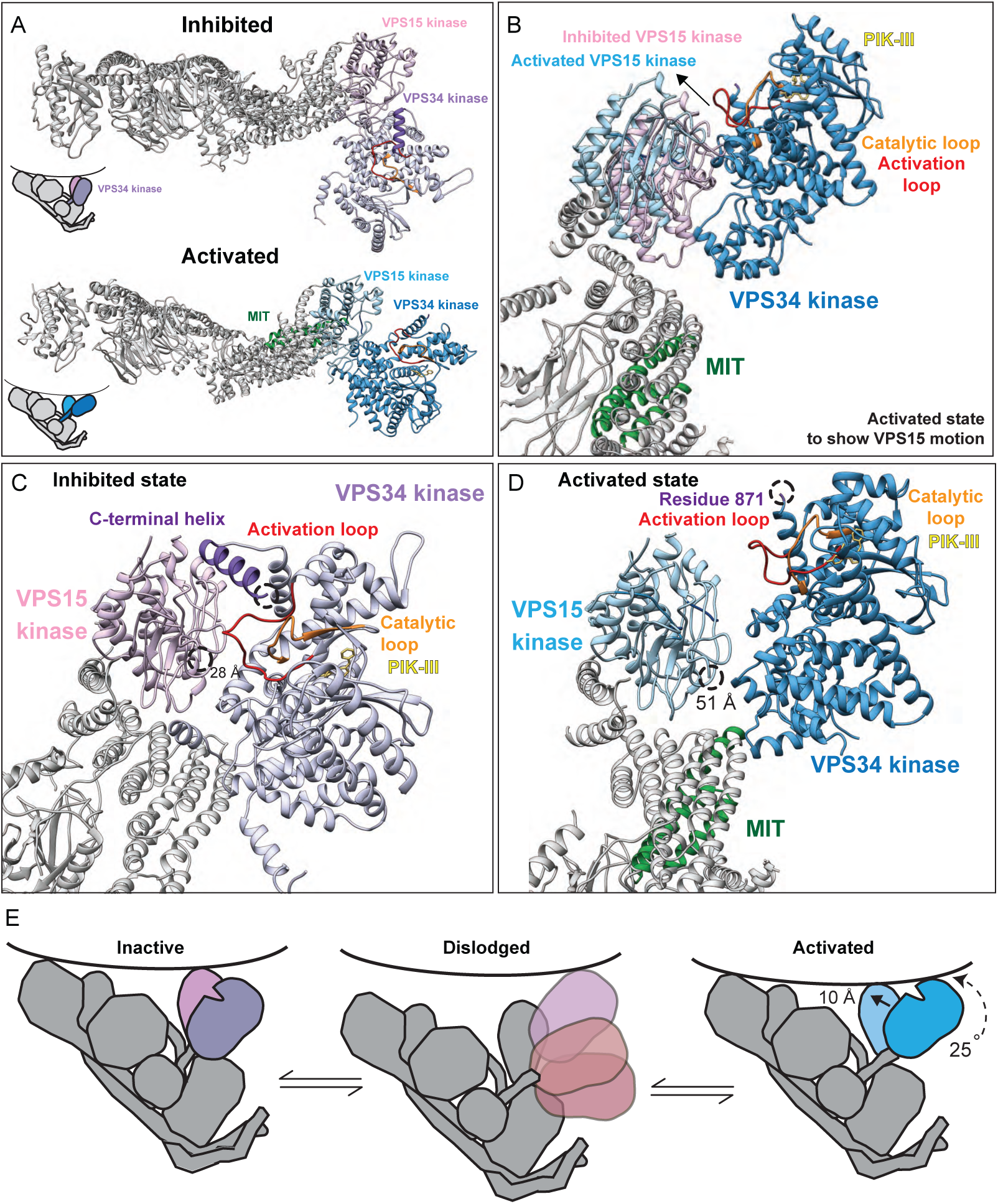
Activation Mechanism for transitioning PI3KC3-C1 from its inhibited to its activated state. (A) Top-down view of the inhibited PI3KC3-C1 (yeast PI3KC3-C2, 5DFZ) state, the VPS34 kinase domain sits perpendicular to VPS15. Top-down view of the activated PI3KC3-C1 (models from 5DFZ, 4PH4, 4ZEY, 4DDP) state docked into the cryoEM reconstruction (EMDB-20390), the VPS15 kinase domain shifts by 10 Å to come more in line with rest of the complex. While VPS34 pivots 25°, the base of the VPS34 kinase domain moves by 5 Å while the top moves by 45 Å. (B) The inhibited state (pink) of VPS15 moves 10 Å away from VPS34 (blue) to reach its activated state (cyan), exposing the activation loop (red) of VPS34. (C) In the inhibited state, the activation loop (red) of VPS34 (lavender, from inhibited state 5DFZ) is occluded by the VPS15 kinase domain (pink, PDB 5DFZ). (D) In the activated state, the activation loop (red) of VPS34 (blue) is released from VPS15 (cyan) inhibition and is more available to engage substrate. (E) Activation model of PI3KC3-C1 on membranes, in the inhibited state, the activation loop of VPS34 is not in a position to engage substrate. In solution, PI3KC3-C1 is highly dynamic, and the VPS34 kinase domain can dislodge (light pink) from the complex to directly bind membranes without the benefit of the rest of the complex. In the activated state, VPS34 is positioned in a precise geometry such that the activation loop of VPS34 is accessible to substrate in the membrane.

## Discussion

The ultimate goal of structural studies of PI3KC3 complex is to understand how they are switched on and off in autophagy by phosphoregulation and by binding of regulatory factors such as Bcl-2, Ambra1, Rubicon, and NRBF2. The structure of one PI3KC3 complex, yeast PI3KC3-C2, has been determined in an inactive conformation (12). The kinase domain of VPS15 keeps basal activity low by restricting the activation loop of VPS34 (12). Our laboratory previously found that the VPS34 kinase domain is highly dynamic (11, 27). We found that “leashing” the VPS34 kinase domain to the VPS15 kinase domain with a 12-residue linker blocked even basal kinase activity (Stjepanovic et al., 2017). This showed that a substantial structural change relative to the published PI3KC3-C2 conformation was essential for any activity, yet the precise nature of the structural change was not clear. The structural change here involves an increase in the VPS34-C to VPS15-N gap from 28 to 51 Å. The increase in distance on activation is great enough that it would be prevented by a 12-amino acid linker. The structural change observed here is thus sufficient to account for the phenotype of the leashed construct.

NRBF2 was chosen as the model allosteric regulator of PI3KC3 in this study because it is one of the well-established protein activators of PI3KC3-C1. By using NRBF2 as a structural probe, it has now been possible to visualize conformational changes likely to be associated with multiple regulatory mechanisms. The base of the complex, near the first MIT binding site, is where a number of post-translational and regulatory signals converge on the autophagic specific PI3KC3-C1 via the regulatory hub, the N-termini of BECN1 and ATG14 (3). This is the site that is modulated by ULK1, AMPK, and Bcl-2, among prominent positive and negative regulators. A single ordered helix is visualized bound to NRBF2^MIT^ which we provisionally assigned as the BECN1 BH3 domain, because the BH3 domain is the portion of BECN1 which is most protected from HDX upon NRBF2 binding. The BH3 is also the binding site for the PI3KC3 inhibitor Bcl-2, suggesting there could be some interplay between regulation by Bcl-2 and NRBF2. The proximity of the BECN1 and ATG14 N-terminal domains to NBRF2 MIT and to the conformationally labile VPS15 solenoid suggest that other N-terminal interactors and post-translational modifications could also act through the NRBF2 binding site and so communicate with the most distant VPS34 lipid kinase domain.

Full length NRBF2 can dimerize of PI3KC3-C1, forming a dimer of pentamers (17, 25), while full length Atg38 does not show this dimerizing effect with respect to yeast PI3KC3-C1 (25). We previously hypothesized that full-length NRBF2 could tether two copies of PI3KC3-C1 within the cell, perhaps facilitating vesicle tethering or membrane binding interactions. The finding that two copies of NRBF2^MIT^ are necessary for full PI3KC3-C1 activation argues that the activating mode of binding is probably one NRBF2 dimer per PI3KC3-C1 complex, which is incompatible with the C1 dimerization model.

NRBF2 (and yeast Atg38) are specific for PI3KC3-C1, and thus specific for autophagy initiation. Our previous HDX-MS work and the cryo-EM work shown here, however, did not show evidence of direct contacts with the C1-unique subunit ATG14. The position of the NRBF2 MIT overlaps with the UVRAG C2 domain in the structure (Fig. S4). This suggests that the presence of the UVRAG C2 domain is a main factor preventing NRBF2 from binding to PI3KC3-C2.

PI3KC3 complex structures have now been determined with the VPS34 catalytic domain in stable active and inactive conformations. The VPS34 catalytic domain is also capable of dislodging from the rest of the complex and sampling a wide range of conformations (11). We speculate that dislodging could facilitate the transition between the active and inactive conformations. The activity of the dislodged state appears to be intermediate between that of the stable inactive and active states (Fig. 4E). As judged by the phenotype of the leashed complex (27), the stable inactive conformation has almost no enzyme activity. The activity of the dislodged state seems likely to reflect that of the isolated VPS34 kinase domain, which is about ∼10% of that of PI3KC3-C1 (27). The presence of a single NRBF2 MIT domain appears sufficient to block the stable inactive conformation, but insufficient to drive full occupancy of the stable active conformation. By contrast, both MIT domains together not only block the stable inactive conformation, they also promote high occupancy of the stable active structure. In this structure, not only are the catalytic site, the C-terminal membrane binding helix, and the activation loop of VPS34 unrestricted, but they are also presented to the membrane substrate with an ideal geometry for PI headgroup phosphorylation.

## Acknowledgments

We thank D. Toso, P. Tobias, P. Grob for cryo-EM support and members of the UC Berkeley cryo-EM supergroup for helpful discussions. This research was supported by NIH grants P01 GM051487 (J.H.H.), R01 GM111730 (J.H.H.), F99 CA223029 (L.N.Y.) and the Bakar Fellows Program (J.H.H.). J.H.H. is a founder of Casma Therapeutics.

## Accession numbers

The EM density maps are being deposited in the EMDB. MIT-Fusion cryoEM map, mask for refinement, and FSC curve is deposited under EMDB-20387. Activated PI3KC3-C1 and full length NRBF2 cryoEM map, the mask used for refinement, and the FSC curve is deposited under EMDB-20390.

## Materials and methods

### Purification of NRBF2 and NRBF2^MIT^

The full length DNA encoding NRBF2 was cloned into vectors 1M (Addgene #29565), generating His6-MBP-TEV-NRBF2. Vectors were obtained from (QB3 Macrolab, UC Berkeley). The NRBF2 truncation construct was subcloned into the vector 1M, generating His6-MBP-TEV-NRBF2^MIT^. BL21 DE3 competent cells were transformed with NRBF2 constructs. Cells were cultured to an OD_600_ of 0.6-0.8 in the presence of kanamycin (0.05 mg/ml) and induced with 0.2 mM IPTG for 3 hours at 37°C. Cell pellets were resuspended in 50 mM HEPES pH 8.0, 200 mM NaCl, 1 mM TCEP, 1 mM PMSF (dissolved in ethanol) and sonicated on ice. Lysates were clarified by centrifugation (18,000g for 60 minutes, 4C). The supernatant was incubated with TALON resin, CloneTech (His6-GFP-TEV-NRBF2) or Amylose resin, New England BioLabs (His6-MBP-TEV-NRBF2^MIT^) for 1.5 hours. Protein was eluted with 300 mM Imidazole (His6-GFP-TEV-NRBF2) or 30 mM Maltose (His6-MBP-TEV-NRBF2^MIT^). His6-MBP-TEV-NRBF2^MIT^ was concentrated with a 10 kDa MWCO Amicon concentrator and injected over a Superdex-200 column (GE Healthcare) equilibrated in 20 mM HEPES pH 8.0, 200 mM NaCl, 2 mM MgCl_2_, 1 mM TCEP.

### Purification of PI3KC3-C1 and MIT-Fusion

The full length DNAs encoding VPS15, VPS34, and BECN1 were codon optimized for expression in HEK293 cells as previously described (11). Full length ATG14 was cloned into a pCAG vector with GST-tag and used for kinase assays. A truncation of ATG14 lacking the last 11 amino acids was similarly cloned into a pCAG vector containing a GST tag (generating GT-ATG14^ΔALPS^) and used for HDX-MS, EM, and SEC studies. Inhibitory phosphorylation sites occur at serines 113 and 120 in NRBF2 (23). These sites were mutated to phosphonulls, creating S113A and S120A. The MIT-linker-BECN1 construct was then generated by amplifying residues 1-159 of NRBF2 including S113A and S120A with an overlapping linker sequence and performing two-step polymerase chain reaction with BECN1 and moved into vector 6A obtained from UC Berkeley macrolab (Addgene # 30124). Cells were lysed by gentle shaking in lysis buffer (50 mM HEPES, pH 8.0, 200 mM NaCl, 10% (vol/vol) glycerol, 1% (vol/vol) Triton X-100, 10 mM TCEP, and protease cocktail of leuptin (final concentration, 10 μM), AEBSF (final, 1mM), and benzamidine (final, 4 mM). Lysates were clarified by centrifugation (18,000×g for 60 min at 4°C) and incubated with glutathione Sepharose 4B (GE Healthcare, Uppsala, Sweden), applied to a gravity column, washed, and purified complexes were eluted with 50 ml wash buffer containing 50 mM reduced glutathione. Eluted complexes were treated with TEV protease at 4°C overnight. TEV-treated complexes were loaded on a 2.5 ml Strep-Tactin Sepharose gravity flow column (IBA GmbH, Göttingen, Germany; at 4°C). The Strep-Tactin Sepharose column was washed, and purified complexes were eluted with wash buffer containing 10 mM desthiobiotin (Sigma-Aldrich, St. Louis, MO). Eluted complexes were purified to homogeneity by injection on Superose 6 16/50 (GE Healthcare) column that was equilibrated in gel filtration buffer (20 mM HEPES, pH 8.0, 200 mM NaCl, 2 mM MgCl_2_, and 1 mM TCEP).

### Electron microscopy sample preparation

Negatively stained samples of PI3KC3-C1 and PI3KC3-C1:MBP-NRBF2^MIT^ were prepared on continuous carbon grids that had been plasma cleaned in a 10% O2 atmosphere for 10 s using a Solarus plasma cleaner (Gatan Inc., Pleasanton, CA). 4 µl of PI3KC3-C1 at a concentration of 25 nM in 20 mM Tris, pH 8.0, 200 mM NaCl, 2 mM MgCl_2_, 1 mM TCEP, and 3% trehalose were placed on the grids and incubated for 30 s. The grids were floated on four successive 50 µl drops of 1% uranyl formate solution incubating for 10 s on each drop. The stained grids were blotted to near dryness with a filter paper and air-dried. MIT-Fusion and MIT-Fusion: MBP-NRBF2^MIT^ samples were imaged using an FEI Tecnai 12 electron microscope (FEI, Hillsboro, OR) operated at 120 keV at a nominal magnification of 49,000 (2.18 Å calibrated pixel size at the specimen level) using a defocus range of −1.5 to −3 µm with an electron dose of 35e^−^/Å^2^. Images were acquired on a TVIPS TemCam F-416 4049 × 4096 pixel CMOS detector (TVIPS GmbH, Gauting, Germany) using the automated Leginon data collection software (28).

### Cryo-EM sample preparation and data acquisition

A sample of MIT-Fusion containing the following subunits: VPS15, VPS34, MIT-12 residue linker-BECN1 fusion, ATG14 (200 nanomolar) was incubated in the presence of BS3 crosslinker for 20 minutes at 1 micromolar concentration. The sample was protected with 0.01% (v/v) NP40 substitute and 2% (w/v) trehalose. The sample incubated for 3 minutes at 5°C, 100% humidity on a Protochips C-flat 2/1 of 300 mesh grid coated with a carbon support, that had been glow-discharged in the presence of amylamine for 55 seconds. The sample was blotted for 3 seconds at blot force 8 and then plunge frozen within Mark IV Vitrobot into 100% ethane. The grid was transferred to a FEI Titan Krios microscope operating at 300 kV and images acquired with a Gatan K2 Direct Electron Detector with a final pixel size of 1.15 Å. Defocus was randomized between −1.3 to −3.3 microns. SerialEM was used to collect an automated dataset of micrographs. A sample of PI3KC3-C1 containing VPS15, VPS34, BECN1, ATG14, and two-fold molar excess NRBF2 (150 nanomolar PI3KC3-C1:300 nanomolar NRBF2) was incubated on ice for 20 minutes in the presence of BS3 crosslinker at 1 micromolar final concentration. The sample was protected with 0.01% (v/v) NP40 substitute and 2% (w/v) trehalose. The sample incubated for 3 minutes at 5°C, 100% humidity on a Protochips C-flat 2/1 of 300 mesh grid coated with a carbon support, and had been glow-discharged in the presence of amylamine for 55 seconds. The sample was blotted for 3 seconds at blot force 8 and then plunge frozen within Mark IV Vitrobot into ethane. The grid was transferred to a FEI Titan Krios microscope operating at 300 kV and images acquired with a Gatan K2 Direct Electron Detector with a final pixel size of 1.15 Å. Defocus was randomized between −1.3 to −3.3 microns. SerialEM was used to collect an automated dataset of micrographs.

### Cryo-EM image processing

One dataset of apo PI3KC3-C1 was collected and processed. Two different datasets of MIT-Fusion were collected, but initially processed separately until 3D refinement. Image stacks were corrected for drift using MotionCor2 (29) and dose-weighted within FOCUS. Contrast transfer functions (CTF) were estimated per micrograph with GCTF and reference-free particle picking was performed on the PI3KC3-C1 dataset and the first dataset of MIT-Fusion using gautomatch (K. Zhang, MRC-LMB, Cambridge). Particles were extracted in Relion-3.0 (30) with box size 352^2^, binned 8-fold. After 2D classification, particles from protein-containing classes were unbinned and used for ab-initio modelling in cryosparc (31). The ab-initio models of PI3KC3-C1 and MIT-Fusion were re-imported, refined in Relion and re-classified in 2D. Upon manual inspection of the severely anisotropic 3D reconstruction, it was decided to stop processing the PI3KC3-C1 dataset. For the MIT-Fusion dataset, six class averages were selected as templates for reference-based particle picking with gautomatch on micrographs from both datasets.

Particles were cleaned up in several rounds of 2D classification, leaving 243,176 and 464,778 particles from the first and second dataset, respectively. The particles from the first dataset were used to build an ab-initio model in Cryosparc-v0.6.5. This model was subsequently refined in Relion using a reference low-pass-filtered to 60 Å, to remove high resolution features which might bias the reconstruction. Masks were created based on the refined map within Relion. Particles were separated into five classes using 3D classification with angular sampling of 7.5° and a mask masking out the kinase domain (Figure S2). The 43,089 particles in the best class were combined with the 464,778 particles from the second dataset. Particles were then separated into three classes using 3D classification without angular sampling and the best class (74,303 particles) was refined. 3D classification was repeated with angular sampling of 7.5° on the refined map using a mask and refinement was performed on the best class (45,236 particles). A dataset of 2NRBF2:PI3KC3-C1 was collected (1700 micrographs) and processed within RELION 3.0.4 (30) using MotionCor2 (29) and GCTF (32) with box size 352^2^, binned 8-fold. After 2D classification, particles from protein-containing classes were unbinned and used as references within RELION for automated particle picking. The resulting 1e6 particles were cleaned up and an ab-initio model was generated in CryoSparc (31). The ab-initio model and the particle stack containing the initial angles from CryoSparc were imported and then refined in RELION first, without a mask (10 Å map) and then with a mask (6.9 Å). Three classification was performed on the 251,100 particles and the particles with the best density for VPS34 were selected. These particles were refined and then subjected to a signal subtraction procedure in which the core solenoid was the region of interest for further alignment, as this is the best region of the map with the least heterogeneity, allowing for an effective signal subtraction step. The core was then subjected to 3D classification without angular sampling and the resulting 86,218 particles were further refined to 6.6 Å and a b-factor of −254 Å^2^.

### CryoEM Modeling

Coordinates were docked into the MIT-Fusion cryoEM reconstruction using homology models from yeast PI3KC3-C2 (PDB: 5DFZ), coordinates from human NRBF2^MIT^ (PDB:4ZEY), and human BECN1 BARA domain (PDB: 4DDP) using UCSF Chimera’s fit Map in Model tool. Swiss-modeler was used to generate the human homology models for VPS15, VPS34, ATG14, and BECN1 using PDB 5DFZ. For the 2NRBF2-PI3KC3-C1 cryoEM reconstruction, coordinates were docked using homology models from yeast PI3KC3-C2 (PDB: 5DFZ), the coordinates of the human NRBF2^MIT^ (PDB:4ZEY), human BECN1 BARA domain (PDB: 4DDP), and the human VPS34 kinase domain (PDB:4PH4) using UCSF Chimera’s fit Map in Model tool. When appropriate, helices were adjusted manually in Coot using real space refinement tools.

### HDX-MS

Amide hydrogen exchange mass spectrometry (HDX-MS) was initiated by a 20-fold dilution of stock PI3KC3-C1 (2 µM), PI3KC3-C1 (2 µM) incubated with full-length NRBF2 (5 µM), or MIT-Fusion (2 µM) into D_2_O buffer containing 20 mM HEPES (pD 8.0), 200 mM NaCl, 2 mM MgCl_2_, and 1 mM TCEP at 30°C. Incubations in deuterated buffer were performed for 10 seconds. Backbone amide exchange was quenched at 0°C by the addition of ice-cold quench buffer (400 mM KH_2_PO_4_ /H_3_PO_4_, pH 2.2). Quenched samples were injected onto a chilled HPLC setup with in-line peptic digestion and desalting steps. Desalted peptides were eluted and directly analyzed by an Orbitrap Discovery mass spectrometer (Thermo Scientific, Waltham, MA). The HPLC system was extensively cleaned between samples. Initial peptide identification was performed via tandem MS/MS experiments. A PEAKS Studio 7 (www.bioinfor.com) search was used for peptide identification. Initial mass analysis of the peptide centroids was performed using HDExaminer version 1.3 (Sierra Analytics, Modesto, CA), followed by manual verification of each peptide. The deuteron content of the peptic peptides covering PI3KC3-C1 was determined from the centroid of the molecular ion isotope envelope. The deuteron content was adjusted for deuteron gain/loss during pepsin digestion and HPLC. Experiments were performed in duplicate.

### Lipid kinase assay

0.78 µmol phosphatidylserine and 0.12 µmol phosphatidylinositol (Avanti Polar Lipids Inc.) were mixed and desiccated overnight. After resuspension in 1 ml buffer (10 mM HEPES pH 8.0, 100 mM NaCl), the lipids were sonicated for 15 min (2 s on/2 s off, amplitude 50%) to generate SUVs. PI3KC3-C1 activity was measured using the ADP-kinase Glo kit (Promega) which converts ADP formation into a luminescence signal. Following pre-incubation of PI3KC3-C1 (22 nM) ± NRBF2 (1.4 µM) with SUVs ([PI] = 20 µM) for 30 min at room temperature, the reaction was started by adding ATP to a final concentration of 50 µM. The kinase reaction had a final volume of 20 µl, buffered with kinase reaction buffer (10 mM HEPES pH 8.0, 100 mM NaCl, 10 mM MgCl_2,_ 1 mM MnCl_2_). After 30 min, the reaction was stopped with 20 µl of ADP Glo reagent (kit) and incubated for another 40 min. The luminescence signal was developed for 20 min after adding 40 µl kinase detection reagent (kit) and measured with a GloMaxMulti plate reader (Promega). The average of three experiments performed in duplicate and normalized to PI3KC3-C1 activity were determined. The error bars represent the standard error of the mean and p-values were determined between PI3KC3-C1 and NRBF2^MIT^ or NRBF2, and MIT-Fusion and MIT-Fusion with NRBF2 using a Tukey comparison.

## Supplementary data

### Supplementary Figure Legends

**Figure S1.**
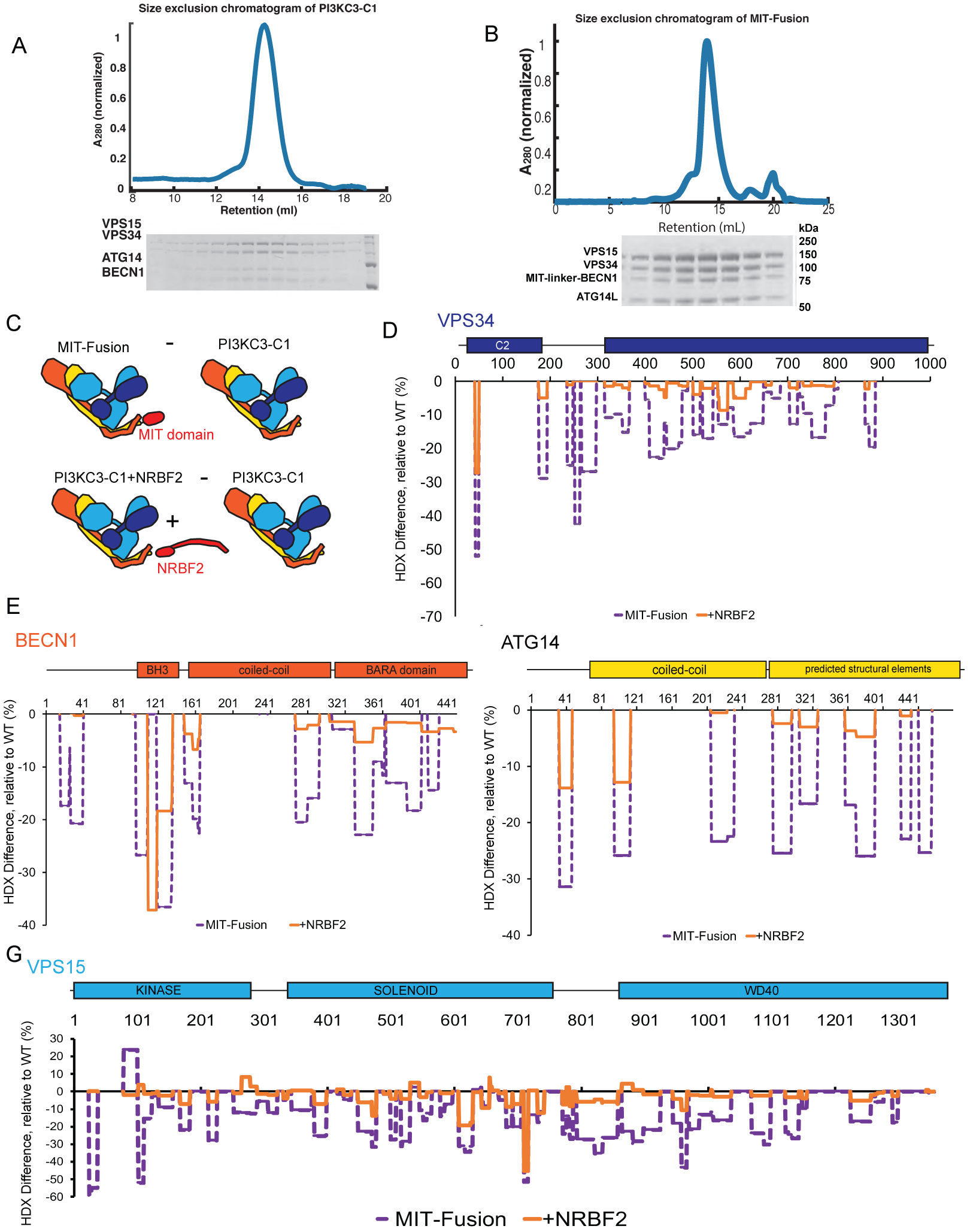
MIT-Fusion sample integrity. Size exclusion profiles and SDS-PAGE of (A) PI3KC3-C1 and (B) BECN1-NRBF2^MIT^ PI3KC3-C1. (C) Schematic of the HDX analysis in which the percent HDX exchange is plotted relative to PI3KC3-C1 for MIT-Fusion (BECN1-NRBF2^MIT^ PI3KC3-C1) and PI3KC3-C1+NRBF2. (D) HD exchange percent for VPS34 (E) HD exchange percent for BECN1 (F) HD exchange percent for ATG14 (G) HD exchange percent for VPS15.

**Figure S2.**
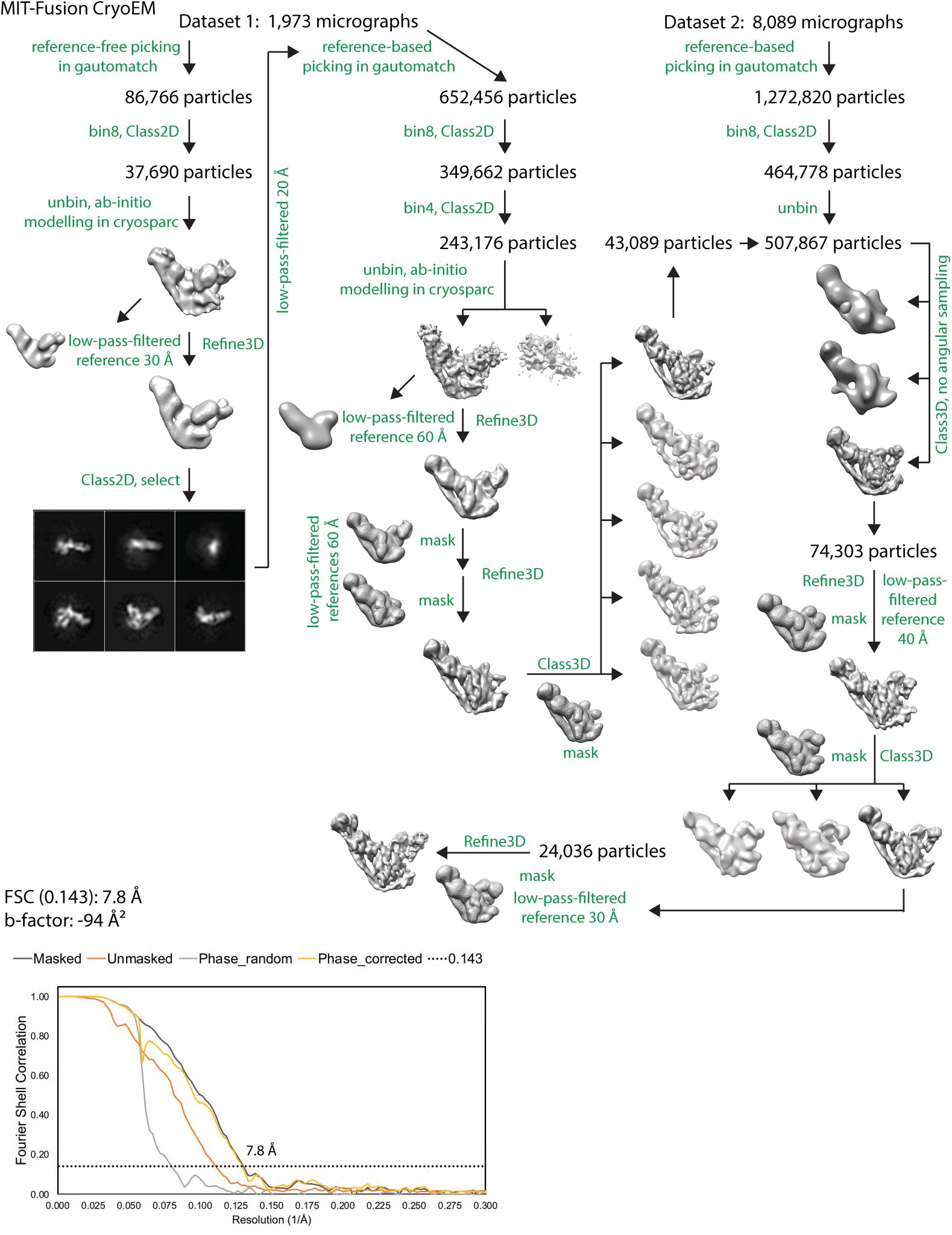
CryoEM on MIT-Fusion. Workflow, FSC curve for MIT-Fusion

**Figure S3.**
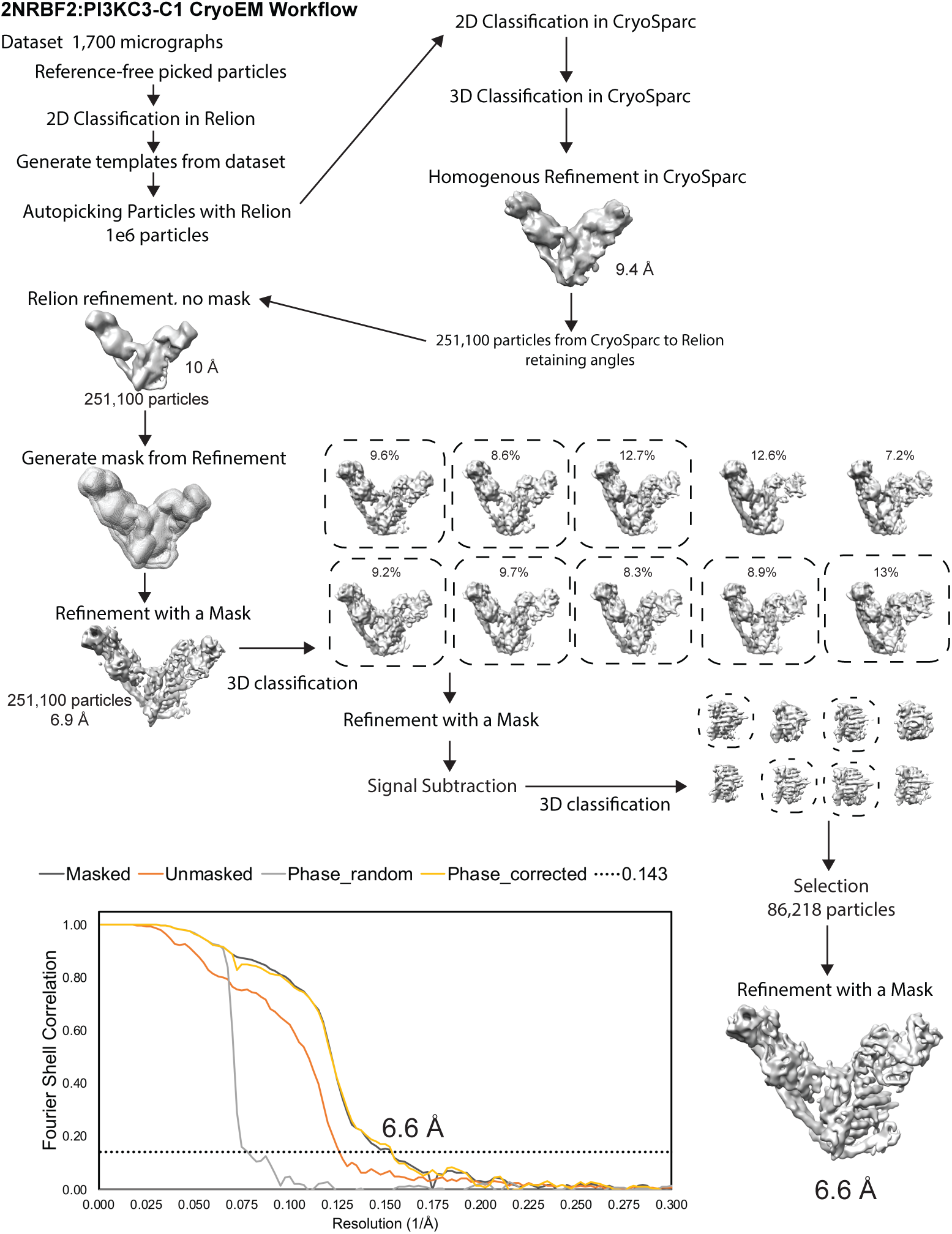
CryoEM on Activated PI3KC3-C1. Workflow, FSC curve for 2NRBF2:PI3KC3-C1 cryoEM map.

**Figure S4.**
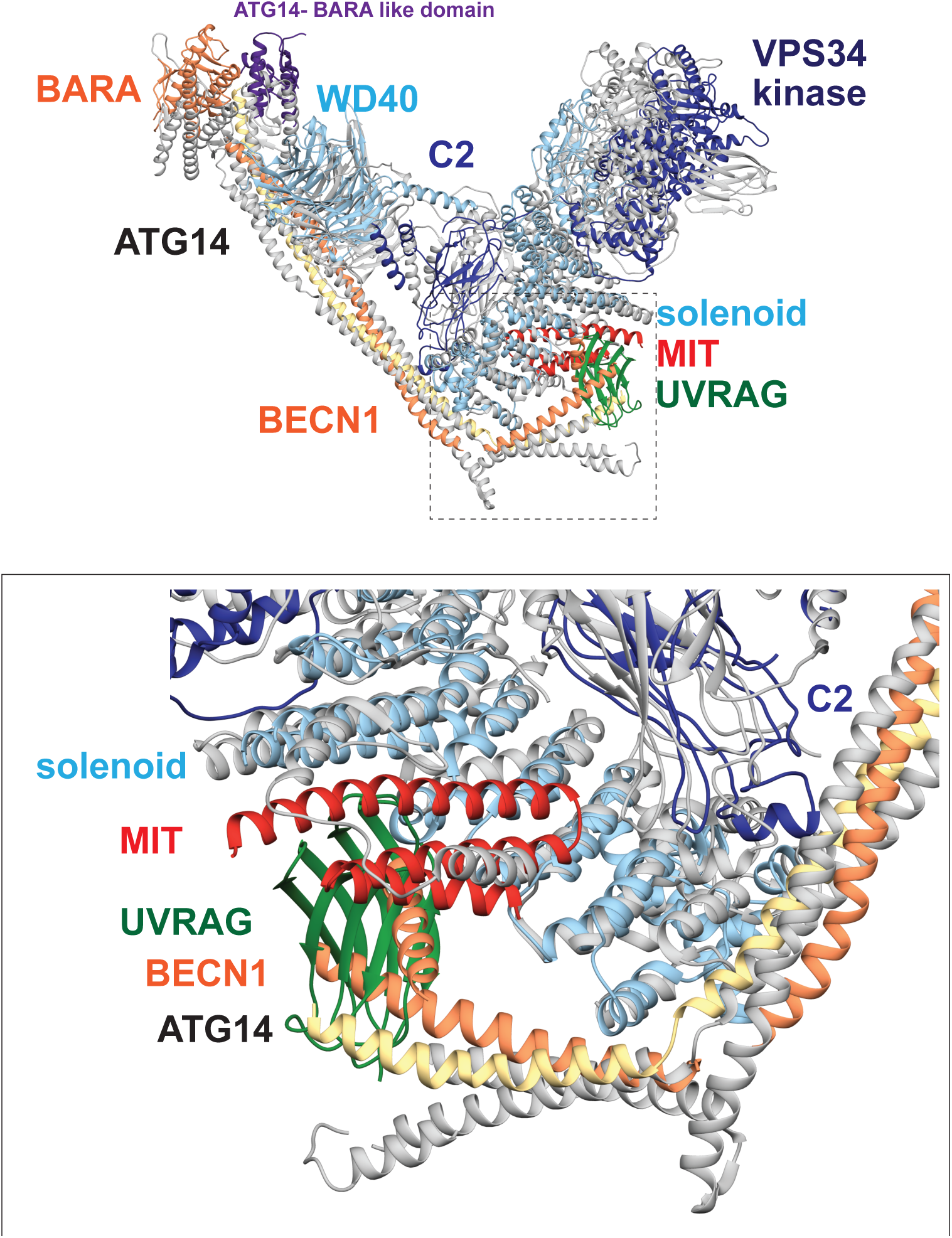
NRBF2 binding site in PI3KC3-C1 vs. Vps38 binding site in PI3KC3-C2.

**Table S1.** Statistics of cryo-EM data collection and processing.

| Map | MITFusion<br>Dataset 1 | MIT-Fusion<br>Dataset 2 | 2NRBF2:PI3KC3-C1 |
| --- | --- | --- | --- |
| **EMDB** | **EMDB-20387** |  | **EMDB-20390** |
| Microscope | Titan Krios | Titan Krios | Talos Arctica |
| Voltage (kV) | 300 | 300 | 200 |
| Detector | K2 Summit | K2 Summit | K3 |
| Recording mode | Super resolution | Super resolution | Super resolution |
| Magnification | 105,000x | 105,000x | 36,000x |
| Movie micrograph pixel size (Å) | 1.15 | 1.15 | 1.14 |
| Dose rate (e-/Å<sup>2</sup>/sec) | 1.52 | 1.22 | 0.938 |
| No. of frames / movie micrograph | 26 frames | 33 frames | 64 frames |
| Frame exposure time | 0.2 sec | 0.2 sec | 0.2 sec |
| Movie exposure time | 5.2 sec | 6.6 sec | 6.4 sec |
| Total dose (e-/Å<sup>2</sup>) | 39.6 | 40.2 | 60 |
| Defocus range (um) | -1.5 to -3 microns | -1.5 to -3 microns | -1.5 to -3 microns |
| EM data processing |  |  |  |
| Number of movie micrographs | 1,973 | 8,098 | 1,700 |
| Box size | 352 | 352 | 352 |
| Particle number (total) | 652,456 | 1,272,820 | 1,004,709 |
| Particle number (post 2D) | 243,176 | 464,778 | 1,004,179 |
| Particle number (post 3D) | 43,098 | 45,236 | 251,100 |
| Particle number (used in final map) | 24,036 |  | 86,218 |
| Refinement angular accuracy (°) | 3.281 |  | 2.704 |
| Refinement offset accuracy (pixels) | 1.729 |  | 1.456 |
| Symmetry | C1 |  | C1 |
| Map resolution, FSC 0.143 | 7.8 Å |  | 6.6 Å |
| Map Resolution Range | 6.5 – 24 Å |  | 5.4 – 18 Å |
| Map sharpening B-factor (Å<sup>2</sup>) | -94 |  | -254 |

## References

1. Levine B, Kroemer G (2019) Biological Functions of Autophagy Genes: A Disease Perspective. Cell 176(1–2):11–42.

2. Backer JM (2016) The intricate regulation and complex functions of the Class III phosphoinositide 3-kinase Vps34. Biochem J 473(15):2251–2271.

3. Hurley JH, Young LN (2017) Mechanisms of autophagy initiation doi:10.1146/annurev-biochem-061516-044820.

4. Ohashi Y, Tremel S, Williams RL (2019) VPS34 complexes from a structural perspective. J Lipid Res 60(2):229–241.

5. Bilanges B, Posor Y, Vanhaesebroeck B (2019) PI3K isoforms in cell signalling and vesicle trafficking. Nat Rev Mol Cell Biol. doi:10.1038/s41580-019-0129-z.

6. Dooley HC, et al. (2014) WIPI2 Links LC3 Conjugation with PI3P, Autophagosome Formation, and Pathogen Clearance by Recruiting Atg12-5-16L1. Mol Cell 55(2):238–252.

7. Galluzzi L, Pedro JMB, Levine B, Green DR, Kroemer G (2017) Pharmacological modulation of autophagy?: therapeutic potential and persisting obstacles. Nat Publ Gr 16(7):487–511.

8. Obara K, Sekito T, Ohsumi Y (2006) Assortment of Phosphatidylinositol 3-Kinase Complexes — Atg14p Directs Association of Complex I to the Pre-autophagosomal Structure in Saccharomyces cerevisiae. 17(April):1527–1539.

9. Itakura E, Kishi C, Inoue K, Mizushima N (2008) Beclin 1 Forms Two Distinct Phosphatidylinositol 3-Kinase Complexes with Mammalian Atg14 and UVRag. Mol Biol Cell 19:5360–5372.

10. Liang C, et al. (2006) Autophagic and tumour suppressor activity of a novel Beclin1-binding protein UVRAG. Nat Cell Biol 8(7):688–99.

11. Baskaran S, et al. (2014) Architecture and dynamics of the autophagic phosphatidylinositol 3-kinase complex. Elife 3. doi:10.7554/eLife.05115.001.

12. Rostislavleva K, et al. (2015) Structure and flexibility of the endosomal Vps34 complex reveals the basis of its function on membranes. Science (80-) 350(6257):aac7365–aac7365.

13. Pattingre S, et al. (2005) Bcl-2 antiapoptotic proteins inhibit Beclin 1-dependent autophagy. Cell 122(6):927–939.

14. Chang C, et al. (2019) Domain Dynamics Article Bidirectional Control of Autophagy by BECN1 BARA Domain Dynamics. 339–353.

15. Cheng X, et al. (2017) Pacer Mediates the Function of Class III PI3K and HOPS Complexes in Autophagosome Maturation by Engaging Stx17. Mol Cell 65(6):1029–1043.e5.

16. Kim Y-M, et al. (2015) mTORC1 Phosphorylates UVRAG to Negatively Regulate Autophagosome and Endosome Maturation. Mol Cell 57(2):207–218.

17. Young LN, Cho K, Lawrence R, Zoncu R, Hurley JH (2016) Dynamics and architecture of the NRBF2-containing phosphatidylinositol 3-kinase complex I of autophagy. Proc Natl Acad Sci 113(29):8224–8229.

18. Behrends C, Sowa ME, Gygi SP, Harper JW (2010) Network organization of the human autophagy system. Nature 466(7302):68–76.

19. Araki Y, et al. (2013) Atg38 is required for autophagy-specific phosphatidylinositol 3-kinase complex integrity. J Cell Biol 203(2):299–313.

20. Cao Y, et al. (2014) NRBF2 regulates macroautophagy as a component of VPS34 complex I. (May).

21. Lu J, et al. (2014) NRBF2 regulates autophagy and prevents liver injury by modulating Atg14L-linked phosphatidylinositol-3 kinase III activity. Nat Commun 5(May):3920.

22. Yang C, et al. (2017) NRBF2 is involved in the autophagic degradation process of APP-CTFs in Alzheimer disease models. Autophagy 13(12):2028–2040.

23. Ma X, et al. (2017) MTORC1-mediated NRBF2 phosphorylation functions as a switch for the class III PtdIns3K and autophagy. 8627(February):1–41.

24. Zhong Y, et al. (2014) Nrbf2 Suppresses Autophagy by Modulating Atg14L-containing Beclin 1-Vps34 Protein Complex Architecture and Reducing Intracellular Phosphatidylinositol-3 Phosphate Levels. J Biol Chem 289(38):3–8.

25. Ohashi Y, et al. (2016) Characterization of Atg38 and NRBF2, a fifth subunit of the autophagic Vps34/PIK3C3 complex. Autophagy 8627(September):00–00.

26. Ma M, et al. (2017) Cryo-EM structure and biochemical analysis reveal the basis of the functional difference between human PI3KC3-C1 and -C2. Cell Res 27(8):989–1001.

27. Stjepanovic G, Baskaran S, Lin MG, Hurley JH (2017) Vps34 Kinase Domain Dynamics Regulate the Autophagic PI 3-Kinase Complex. Mol Cell 67(3):528-534.e3.

28. Suloway C, et al. (2005) Automated molecular microscopy: The new Leginon system. J Struct Biol 151(1):41–60.

29. Zheng SQ, et al. (2017) MotionCor2 - anisotropic correction of beam-induced motion for improved cryo-electron microscopy. Nat Methods:4–9.

30. Scheres SHW (2012) RELION: Implementation of a Bayesian approach to cryo-EM structure determination. J Struct Biol 180(3):519–530.

31. Punjanji A, Rubinstein JL, Fleet DA, Brubaker MA (2017) cryoSPARC: algorithms for rapid unsupervised cryo-EM structure determination. Nat Methods In Press(December 2016). doi:10.1038/nmeth.4169.

32. Zhang K (2016) Gctf: Real-time CTF determination and correction. J Struct Biol 193(1):1–12.

